# A novel self-organizing human pluripotent stem cell system reveals the role of Wnt signaling parameters in anterior-posterior patterning of the nervous system

**DOI:** 10.1101/2025.06.07.658391

**Authors:** Siqi Du, Eleni Anastasia Rizou, Aryeh Warmflash

## Abstract

A Wnt activity gradient is essential for the formation of the anterior-posterior (AP) axis in all vertebrates. The relationship between the dynamics of Wnt signaling and specification of AP coordinates is difficult to study in mammalian embryos due to the inaccessibility of developing embryos and the difficulty of live imaging. Here, we developed an *in vitro* model of human neuroectoderm patterning, where the AP axis self-organizes along the radius of a micropatterned human pluripotent stem cell colony. We used this system to study the quantitative relationship between Wnt signaling in space and time and the resulting AP patterns. We found that rather than a smoothly varying gradient along the axis, signaling is elevated in midbrain compared to either surrounding region. The timing, rather than the amplitude or duration, of the Wnt response played the most important role in setting axial coordinates. Loss of WNT8A reduces posterior Wnt activity and anteriorizes cell fates, demonstrating that even cells exposed to high exogenous Wnt require endogenous ligand production to fully posteriorize and to position the midbrain-hindbrain boundary. These results establish a simple system for studying the patterning of the human nervous system and elucidate how cells interpret Wnt dynamics to determine their position along the AP axis.

## 1 Introduction

Establishing the anterior-posterior (AP) axis is a fundamental step in defining the body plan of all vertebrates [1]. The central nervous system originates from the ectodermal germ layer and is compartmentalized along this axis into the forebrain, midbrain, hindbrain, and spinal cord. To establish the AP pattern within the neuroectoderm, pluripotent cells from the epiblast must both be converted to neural fates and assigned axial coordinates. The experiments of Spemann and Mangold demonstrated the existence of an organizer in amphibians capable of converting cells to neural fates within the ectoderm [2, 3]. This neuralizing ability results from secreted inhibitors, and inhibition of both branches of the TGF*ω* pathway—Activin/Nodal and BMP—is sufficient to convert cells to anterior neural fates in multiple vertebrate models, including human pluripotent stem cells (hPSCs) [4–11]. This suggests that the anterior fate is the default, and posteriorizing signals are required to establish the axis, consistent with the activation-transformation model originally proposed by Nieuwkoop [12, 13]. More recently, lineage tracing experiments as well as measurements of chromatin accessibility have suggested that cells in the posterior region acquire regional identity before neural identity and cells of the spinal cord share a common lineage with the somitic mesoderm rather than with more anterior neural fates [14–16].

The molecular identity of the caudalizing signal has been the subject of extensive work. Many candidates have been proposed, including Wnts, FGFs, and retinoic acid (RA) [17]. The Wnt pathway has the ability to affect the entire AP axis without interfering with neural induction [18–21]. Multiple Wnt ligands are expressed in the posterior region of the embryo [22–27]. Meanwhile, Wnt antagonists are found in the anterior region and are critical for head induction [28–33]. Beyond the expression patterns of Wnts and their inhibitors along the AP axis, functionally, an endogenous activity gradient of Wnt/*ω*-catenin also aligns with the AP axis and is indispensable for proper patterning [19, 30, 34]. Additionally, both mouse Wnt reporter lines and nuclear *ω*-catenin staining in Xenopus have revealed local enrichment of Wnt activity in the midbrain, immediately anterior to the midbrain-hindbrain boundary (MHB) [35, 36], pointing to additional spatial structure beyond a simple monotonic gradient.

Stem cell models provide powerful tools for studying mammalian embryos, which are otherwise difficult to access *in vivo* [37, 38]. Beyond differentiating into individual cell types, pluripotent stem cells can recapitulate aspects of embryonic patterning by generating ordered arrangements of multiple fates. For neural development, dual-SMAD inhibition efficiently converts human pluripotent stem cells to neural fates [11], providing a starting point for modeling regional identity along the neural axis. Such protocols have been combined with microfluidics to create bioengineered models of the developing neural tube that successfully recapitulate the cell fates along the AP axis [39, 40]. In these models, the gradients used to establish the axes are externally imposed and so they do not mimic the self-organization of gradients which occurs *in vivo*. We sought to create a self-organizing *in vitro* model system to study patterning along the AP axis with minimal external spatial cues.

Our previous work and that of others showed that globally applying pathway activators or inhibitors to geometrically confined hPSC cultures can recapitulate germ layer or ectodermal mediolateral patterning depending on the protocol (Fig. 1A) [41–47]. These micropatterned systems generate dynamic signaling patterns starting from the edge [48, 49], and we reasoned that self-organized Wnt activity patterns within the neuroectoderm could be generated in a similar manner. In this study, we show that treating colonies differentiating to neural fates with Wnt ligands creates self-organized patterns of cell fates corresponding to the AP axis within the neuroectoderm. Comparison with singlecell RNA sequencing data from mouse embryos [50] revealed that this system successfully replicates the spatial and temporal emergence of cell fates along the AP axis. We used this system to quantitatively dissect the relationship between Wnt signaling dynamics and cell fates and found that the timing of Wnt signaling, rather than the magnitude or duration, defines the boundary between the midbrain and hindbrain. Moreover, removing WNT8A diminishes Wnt activity in the posterior region and shifts cell identities anteriorly across the colony, indicating that endogenously produced ligands are needed both to specify posterior fates and to position the MHB. Our system also revealed that, rather than a monotonic increase in Wnt activity from anterior to posterior, the midbrain region has higher Wnt activity than either of the neighboring territories. These findings highlight the power of self-organizing systems to dissect both the development and interpretation of signaling patterns along the AP axis.

**Figure 1:**
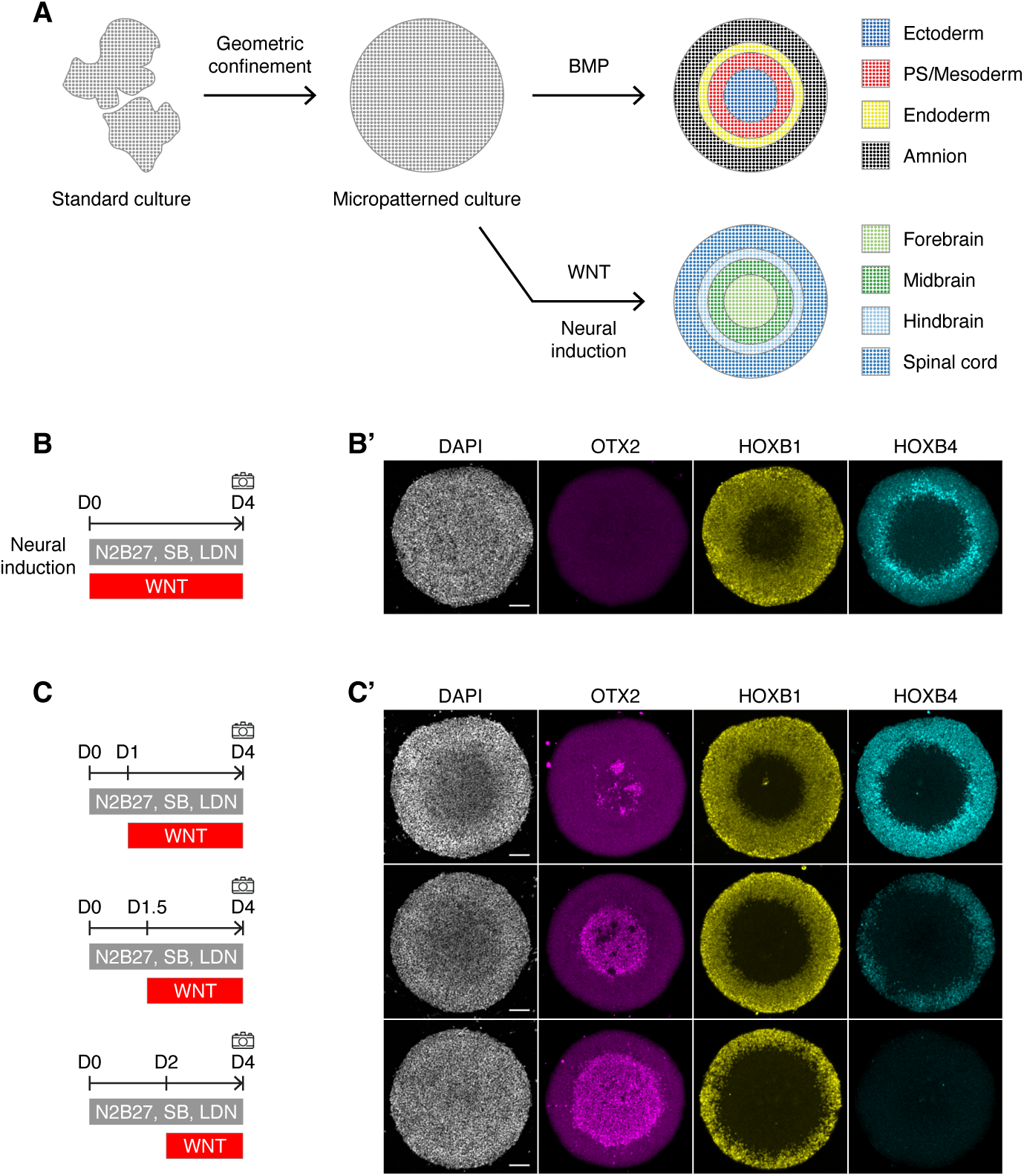
A novel self-organizing system to study anterior-posterior patterning. (A) Schematic comparing protocols for germ layer and neural AP axis micropatterns. (B, C) Diagram of treatments. The camera icon denotes when cells were fixed and observed. (B’, C’) Representative immunofluorescence images of DAPI, OTX2, HOXB1, and HOXB4 following 4 days of neural induction in which 300 ng*/*mL WNT3A was added throughout the differentiation period (B) or at the indicated time points (C). The colony diameter in B’–C’ is 700 µm. Scale bar, 100 µm. Maximum intensity projection images are shown.

## 2 Results

### A micropatterned model of AP patterning within the nervous system

We sought to create a model in which as much of the AP axis as possible, ranging from forebrain to spinal cord, is recapitulated along the radius of a human pluripotent stem cell (hPSC) colony. Based on earlier work with micropatterns, we anticipated that added Wnt ligands would signal most strongly at the colony edge, conferring posterior identities there while allowing more anterior fates to develop at the center (Fig. 1A). Therefore, we performed a neural induction protocol on micropatterns and treated the colonies with WNT3A to initiate AP axis patterning (Fig. 1B). We used OTX2 to mark anterior cell fates, and HOXB1 and HOXB4 to mark posterior cell fates. Under WNT3A treatment, the colony forms concentric rings of HOXB1 and HOXB4, with HOXB1 extending further toward the colony center (Fig. 1B’). This confirms that exogenous Wnt ligands posteriorize the colony edge more strongly than the center, generating cell fates that are spatially organized along the AP axis. The absence of OTX2, however, indicates that this protocol fails to generate the most anterior cell fates, including forebrain and midbrain. We hypothesized that delaying the addition of Wnt might allow some cells to commit to anterior fates, preventing the posteriorization of the entire colony upon ligand stimulation. To test this, we added WNT3A after 1, 1.5, or 2 days of neural differentiation and examined cell fates after 4 days (Fig. 1C–C’). When Wnt was added after 1 day, small, irregular patches of cells expressed OTX2 near the colony center. When addition was delayed by 1.5 days, a circular domain of cells expressed OTX2 in the center with rings of HOXB1 and HOXB4 expression near the edge. If addition was further delayed to 2 days, the domain of OTX2 expanded further but HOXB4 disappeared. Thus, later Wnt addition produces progressively more anterior patterns. We found that addition of WNT3A at day (D) 1.5 is optimal for obtaining the broadest range of cell fates along the AP axis.

### The model recapitulates the dynamics of gene expression that occur *in vivo*

Having demonstrated that the micropatterned system generates cell fates corresponding to a broad range of the AP axis, we next asked how accurately it reflects the dynamics of gene expression *in vivo* across a larger set of markers for specific regions of the nervous system. We obtained a publicly available single-cell (sc) RNA sequencing dataset of mouse embryos collected from embryonic day (E) 6.5 to E8.5, a period that spans the transition of the epiblast into patterned neuroectoderm and corresponds to the processes occurring in the micropatterned culture system [50]. We selected five cell fate groups representing the neuroectoderm and its precursors—rostral neuroectoderm, caudal neuroectoderm, forebrain/midbrain/hindbrain, spinal cord, and neuromesodermal progenitors (NMPs)—and performed scVI normalization and UMAP visualization [51]. This produced a clear ordering of cells along space on the first UMAP coordinate and along time on the second (Fig. 2A, C).

**Figure 2:**
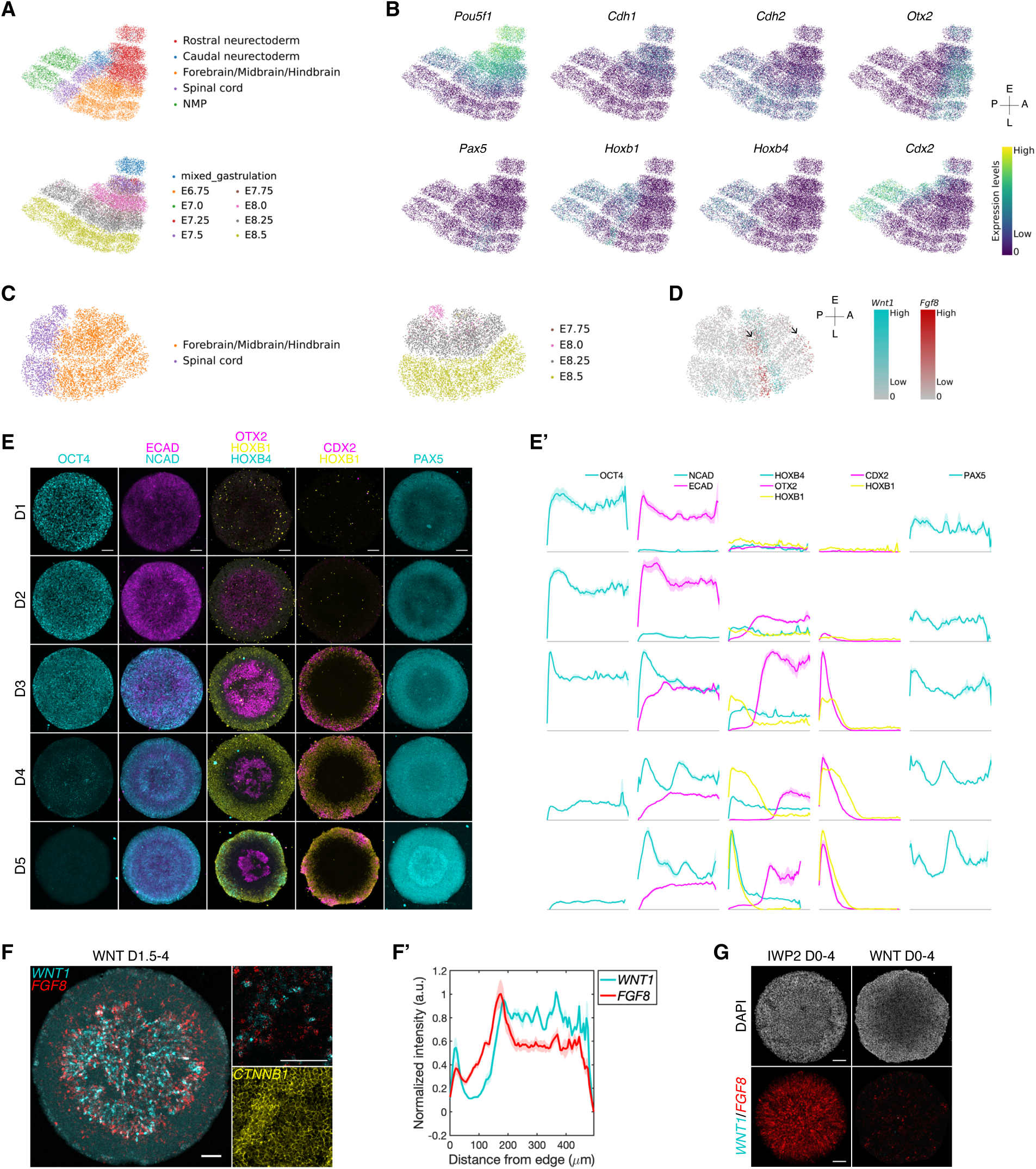
The micropatterned system recapitulates the dynamics of gene expression in mammalian embryos. (A) Uniform manifold approximation and projection (UMAP) plots display 14,937 selected cells from the mouse atlas dataset. Cells are colored by their cell-type (top) or developmental stage (bottom). (B) Expression levels of the indicated genes overlaid on the UMAP plot shown in A. (C) UMAP plots display 6,610 selected cells from the mouse atlas dataset. Cells are colored by their cell-type (left) or developmental stage (right). (D) Expression levels of *Wnt1* (cyan) and *Fgf8* (red) overlaid on the UMAP plot shown in C. (E) Representative immunofluorescence images of the indicated markers from D1–D5. WNT3A was added at D1.5 during the 5-day neural induction. (E’) Quantification of the expression levels of the indicated markers as a function of distance from the colony edge on each day from D1– D5. (F) Representative images of *WNT1* and *FGF8* mRNA expression, along with immunofluorescence for CTNNB1 (*ω*-catenin). The zoomed-in images represent a single z-plane. (F’) Quantification of the expression levels of *WNT1* and *FGF8*, normalized to DAPI intensity. (G) Representative images of *WNT1* and *FGF8* mRNA expression, along with DAPI staining. 4 µM IWP2 or 300 ng*/*mL WNT3A was added during the 4-day neural induction. The colony diameter in F is 1000 µm while in others it is 700 µm. Scale bar, 100 µm.

Temporally, cells first express the pluripotency marker *Pou5f1* (*Oct4*) along with *Cdh1* (Ecad) (Fig. 2B). Expression of these genes decreases as *Cdh2* (Ncad), an early indicator of neural differentiation, increases, with this transition occurring slightly earlier anteriorly than posteriorly. Spatially, from anterior to posterior (right to left on the UMAP), the forebrain marker *Foxg1* partially overlaps with the anterior marker *Otx2*, followed by *Pax5* near the caudal midbrain/rostral hindbrain region [52], and then the posterior markers *Hoxb1*, *Hoxb4*, and *Cdx2* (Fig. 2B, Fig. S1B). *Otx2* is expressed during gastrulation and later becomes restricted to the anterior neuroectoderm. *Pax5*, *Pax6*, and *Foxg1* are expressed at later stages, with *Foxg1* beginning only at E8.5. In the posterior, *Cdx2* expression precedes that of the Hox genes, and *Hoxb1* precedes *Hoxb4*. The boundary between midbrain and hindbrain is a crucial signaling center known as the isthmic organizer, marked by a sharp boundary in *Otx2* expression [53, 54]; the midbrain marker *Wnt1* and the isthmic organizer marker *Fgf8* are expressed in adjacent stripes near the midbrain-hindbrain boundary (MHB) [54, 55], trends also present in the scRNA dataset (Fig. 2D). *Fgf8* expression was additionally found in the forebrain and tailbud (Fig. S1B), consistent with previous literature [56, 57].

For comparison, we performed immunostaining of the same markers in our *in vitro* system on each day from D1 to D5 (Fig. 2E–E’, Fig. S1D–D’). OCT4 is downregulated after D3, and NANOG is expressed only on D1. SOX2 is continuously expressed throughout the colony, with levels that vary by radial position. The switch from ECAD to NCAD occurs on D3, though a belt of lower NCAD expression remains midway through the colony. Among the AP identity markers, OTX2 emerges first on D2, followed by HOXB1 and CDX2 on D3, HOXB4 on D4, and PAX5 between D4 and D5. Thus, the *in vitro* system not only recapitulates the spatial ordering of markers from anterior to posterior but also reproduces the dynamics of cell fate decisions. The temporal correspondence is approximately: D1 to E6.5, D2 to E6.75–E7.5, D3 to E7.5–E8.0, D4 to E8.0–E8.25, and D5–D9 to E8.5 and beyond.

We did not detect markers of the most anterior fates under any of the protocols above. Given that Wnt activity is inhibited in the forebrain [19], we tested whether adding IWP2, a PORCN inhibitor that blocks secretion of all Wnt proteins, would induce forebrain differentiation (Fig. S1C–C’). Low levels of PAX6 and *FGF8* expression were detected on D4, while FOXG1 expression was observed only on D8, indicating that PAX6 and *FGF8* expression precede FOXG1 (Fig. 2G, Fig. S1C). The extended culture time is consistent with the longer maturation of these fates in mouse, particularly for Foxg1-expressing cells [58]. Altogether, these trends agree with the *in vivo* mouse scRNA dataset and the literature on the expression patterns of these genes [58–61].

To study MHB formation in our system, we performed fluorescence *in situ* hybridization for these genes and found that their adjacent localization was well preserved (Fig. 2F–F’). When colonies were treated with IWP2 during neural induction, *WNT1* expression was lost but *FGF8*-expressing cells filled the colony, consistent with FGF8, but not WNT1, expression in the forebrain (Fig. 2G) [56]. Both markers were lost when Wnt ligands were added on D0 rather than D1.5, consistent with a more posterior identity throughout the colony [57]. Taken together, our results show that the micropatterned model produces patterns that faithfully recapitulate a wide range of fates along the AP axis, and that this range can be shifted anteriorly or posteriorly by changing Wnt activity levels or timing.

### Quantitative relationship between Wnt parameters and AP pattern

Having established that Wnt activity is a crucial parameter for controlling AP patterning in our system, as it is *in vivo*, we examined how the parameters of Wnt stimulation influence the resulting patterns by varying the concentration, duration, and timing of WNT stimulation and activity. In the following sections, we focus on WNT3A-treated samples, rather than those that are untreated or IWP2-treated, due to their broader coverage of the AP axis (Fig. 3, Fig. S2). We first varied WNT3A concentration from 0 to 1000 ng/mL, with addition on D1.5 as above. As concentration increased, the OTX2 and PAX5 domains shrank and disappeared once Wnt exceeded 300 ng/mL, while the HOXB1 and HOXB4 domains expanded from the edges and their expression levels rose (Fig. 3A–A’, Fig. S2A–A’). Thus, higher Wnt concentrations produce more posterior patterning in a graded fashion. We next tested the timing of Wnt addition by treating colonies with WNT3A for 24 hours starting on D0, D1, D2, or D3 (Fig. 3B). The MHB, represented by the gap between OTX2 and HOXB1, shifted dramatically with timing: later addition pushed the boundary outward so that anterior fates filled a larger fraction of the colony. When 100 ng/mL WNT3A was added on D1.5, the colonies have a large OTX2-positive domain with no HOXB4 expression, indicating a relatively anterior identity. In contrast, when WNT3A was added on D0, the colonies completely lost OTX2 expression and showed robust HOXB4 expression. These results suggest that while timing and concentration of Wnt are both important parameters, reducing the Wnt concentration does not fully compensate for the posteriorizing effects of early treatment (Fig. S2E).

**Figure 3:**
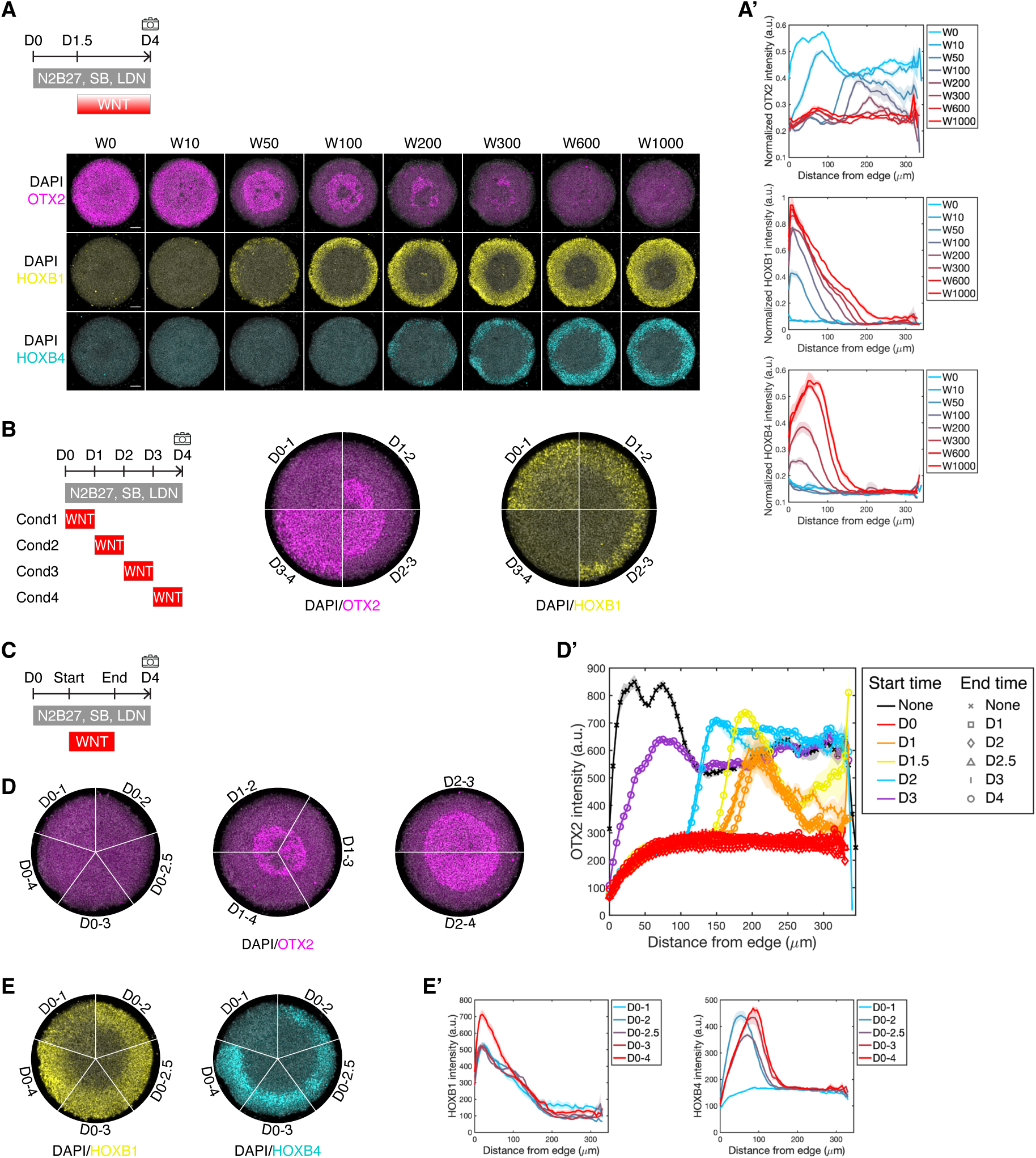
Timing is more critical than duration or concentration in cell fate decisions. (A) Representative immunofluorescence images of the indicated markers. The indicated concentrations of WNT3A (ng*/*mL) were added on D1.5. (A’) Quantification of the expression levels of OTX2, HOXB1, and HOXB4 following the treatments described in A, normalized to DAPI intensity. (B) Representative immunofluorescence images of OTX2 and HOXB1. During the 4-day neural induction, WNT3A was added for 24 hours, at D0, D1, D2, or D3. (C) Schematic of the protocol shows cells treated with neural induction cocktail for 4 days, with varying start and end times of WNT3A, resulting in different durations. (D, E) Representative immunofluorescence images of indicated markers, grouped by the start time of WNT3A treatment. (D’) Quantification of the expression levels of OTX2. The lines are colored by the start time of Wnt treatment, and the marker shapes correspond to the end time. (E’) Quantification of the expression levels of HOXB1 and HOXB4.

Finally, we examined the effect of varying the duration of Wnt exposure by adding WNT3A on a fixed day and removing it on different days. Surprisingly, OTX2 expression was determined solely by the timing of Wnt addition, not by duration (Fig. 3D–D’). When Wnt was added on D0, no OTX2 was expressed even if Wnt was present for only one day; when added on D1, OTX2 remained in the center; when added on D2, OTX2 occupied a much larger territory. The data are summarized in Fig. 3D’, where curves group by starting time rather than ending time or duration. While duration did not shift the position of the anterior-posterior boundary, the posterior cells themselves became progressively more posterior with longer WNT3A exposure (Fig. 3E–E’). This is particularly evident for HOXB4 under D0 treatment, where anterior fates do not form: with increasing duration, the HOXB4 peak shifts inward, reflecting deeper posteriorization rather than movement of the boundary. Thus, among the three parameters, timing is the strongest determinant of the range of anterior fates, while concentration and duration have graded effects, with higher values of either producing more posterior fates.

### Wnt signaling at different times has similar dynamics but different outcomes

To better understand how Wnt parameters influence AP patterning, we measured Wnt signaling activity using H9 cells expressing GFP fused to histone H2B under the control of a TCF/LEF1-responsive promoter [62]. These cells differentiated with AP fate patterns similar to the ESI017 cells under the micropatterned protocols described above (Fig. S3A–B’, E–E’, Fig. S2B–D, Fig. 3A). At the endpoint, however, the patterns of Wnt activity and cell fate did not correspond: for example, treatment from D0 to D1 produced the strongest posteriorization but the lowest endpoint Wnt signaling (Fig. S3A–A’). This motivated us to examine the signaling dynamics rather than only endpoint levels.

We monitored Wnt signaling while exposing cells to WNT3A for 24 hours, varying the day of addition. WNT3A triggered an expanding front of Wnt activity that originated at the colony edge and traveled inward, reaching approximately 100 µm into the colony (Fig. 4A). Cells responded rapidly upon Wnt addition and reporter levels peaked between 24 and 48 hours before declining somewhat thereafter, though signaling remained elevated throughout differentiation (Fig. 4C). Although the response amplitude varied with the day of Wnt addition, the overall dynamics remained consistent (Fig. 4C, right). We then varied the duration of Wnt exposure: longer exposure produced a stronger and more prolonged response, though the response still peaked at approximately 24 hours before declining (Fig. 4B, D). Notably, the signaling wave did not extend beyond 100 µm from the colony edge under any timing or duration we tested; only increasing WNT3A concentration allowed the wave to reach further inward (Fig. S3C–D).

**Figure 4:**
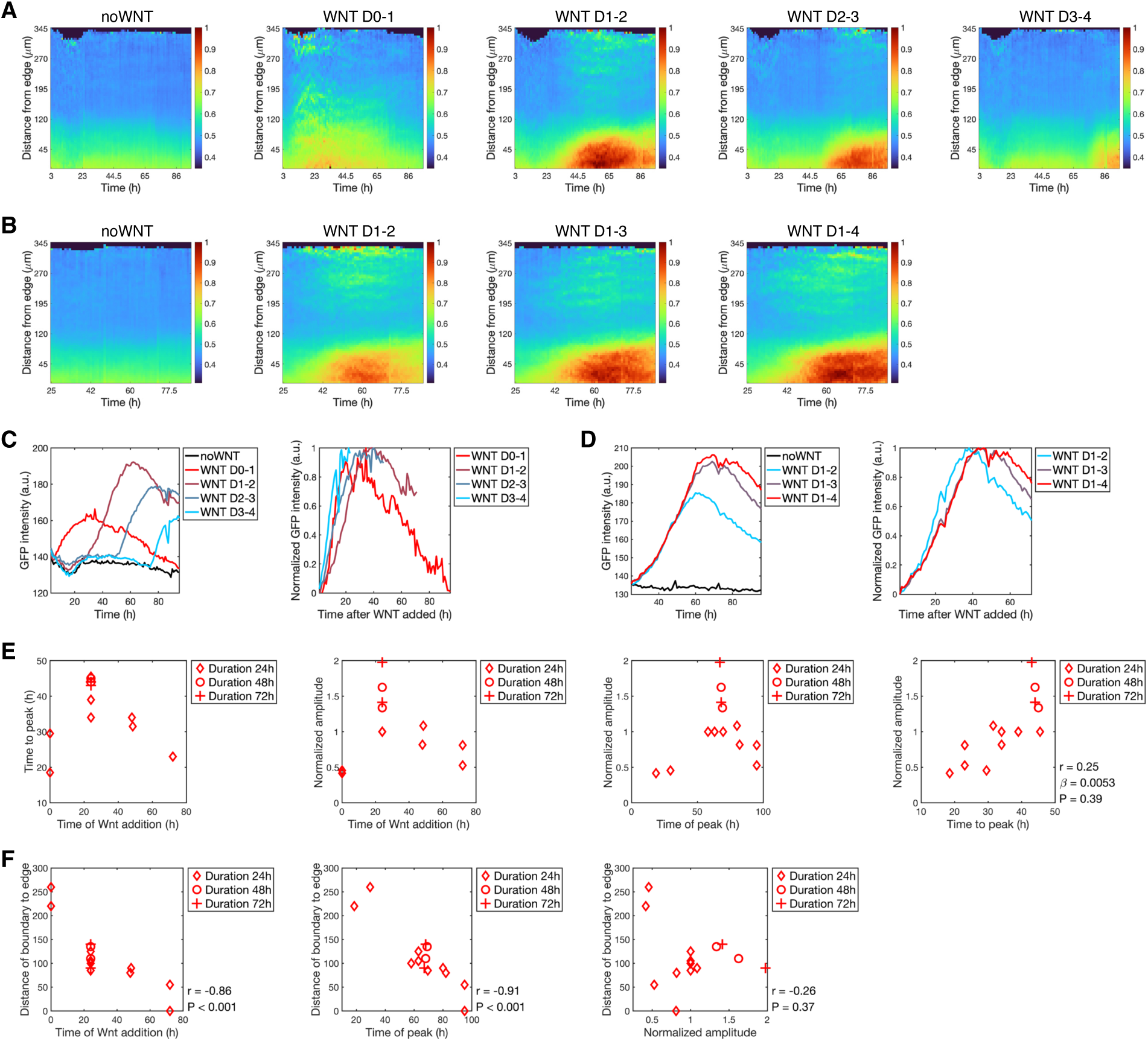
The timing of Wnt signaling determines the location of the OTX2 expression boundary. (A, B) Heatmap plots of GFP levels, with the x-axis representing experimental time and the y-axis representing the distance from the colony edge, following 4 days of treatment with neural induction cocktail and WNT3A as described in the labels. The colony diameter is 700 µm. (C, D) Quantification of average GFP intensity in the ring-shaped domain (0–100 µm from the edge) as a function of time. Normalized GFP intensity is calculated by scaling the GFP levels to their starting (base) and peak (maximum) values. (E, F) The scatter plots combine experiments testing different Wnt durations and timings, with the marker shape indicating the duration. The time of Wnt addition is the time, measured from the onset of differentiation, at which WNT3A was added; the time of peak is the time, on the same scale, at which the Wnt reporter response reaches its maximum; and the time to peak is the interval between the two. Normalized amplitude is the amplitude of the Wnt response rescaled so that the no-Wnt condition is 0 and the WNT D1–D2 condition is 1. The distance from the boundary to the edge is quantified by summing the intensities of OTX2 and HOXB1, and measuring the distance from the edge to the dip in the summed intensity. r is the Pearson correlation coefficient between the two variables on the xand y-axes. P is the p-value. *ω* is the slope of the linear fit.

We examined the relationship between the timing of Wnt addition and both the response amplitude and peak time across all live imaging experiments in which the duration and timing of Wnt exposure were varied. The response was strongest when Wnt was added on D1, with weaker responses when added earlier or later (Fig. 4E, middle left), suggesting a temporal window during neural differentiation when cells are most responsive to Wnt. Because the shape of the response curve remained consistent across treatments (Fig. 4E, right), the response can be summarized by two parameters: peak amplitude and peak timing. We plotted the distance of the HOXB1/OTX2 boundary from the edge as a function of these signaling parameters (Fig. 4F). Later Wnt addition shifted the boundary outward with a linear dependence on timing, whereas peak amplitude showed no clear relationship to boundary position. We conclude that the timing of Wnt signaling is a key determinant of MHB location.

### AP pattern partially scales with colony size

Previous work on gastrulation-stage micropatterns showed that the scale of the pattern remains constant as colony size varies [41]; consequently, when colonies become sufficiently small, the central cell fates are lost. An alternative is scaling, in which the pattern adjusts to match the available space within the tissue or colony. To investigate which regime governs AP patterning, we grew hPSCs on micropatterned colonies with radii ranging from 50 to 500 µm

(Fig. 5A–A’). In the largest colonies, the HOXB1 ring is approximately 150 µm wide; if the pattern scale were invariant, HOXB1 would cover the entire colony and OTX2 would be lost for radii below this size. Instead, colonies of radius 100 µm still display a ring-shaped HOXB1 domain, indicating that this territory shrinks with colony size. However, OTX2 is entirely lost in colonies of radius 50 and 100 µm, showing that scaling is only partial rather than fully preserving the pattern. Consistent with this, at radius 250 µm the OTX2 peak shifts slightly outward compared to larger colonies, partially compensating for the reduced territory.

**Figure 5:**
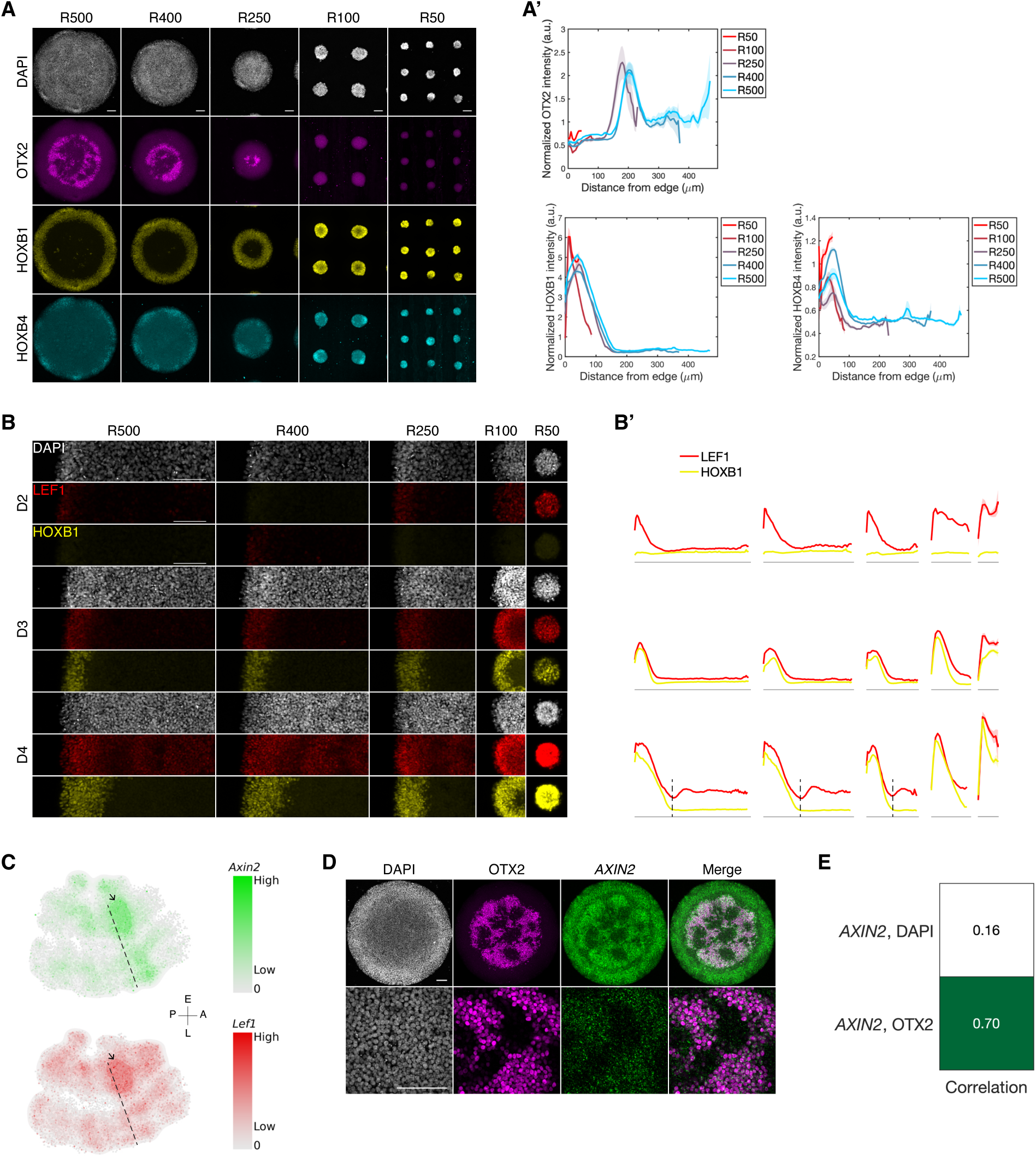
The midbrain-hindbrain boundary corresponds to a local minimum in Wnt signaling. (A) Representative immunofluorescence images of DAPI, OTX2, HOXB1 and HOXB4. The colonies vary in radius, ranging from 50 µm to 500 µm. (A’) Quantification of the expression levels of OTX2, HOXB1 and HOXB4, normalized to DAPI intensity. (B) Cropped representative immunofluorescence images of DAPI, LEF1 and HOXB1 at D2, D3 and D4. (B’) Quantification of the expression levels of the indicated markers as a function of distance from the colony edge on each day from D2–D4. The dashed lines indicate the corresponding HOXB1 levels at the point where LEF1 expression dips. For the indicated colony radii, *n* = 5 for R500, R400 and R250; *n* = 8 for R100; and *n* = 9 for R50. (C) Kernel density estimate (KDE) plots show the estimated probability densities of *Axin2* and *Lef1* expression overlaid on the UMAP plot from Fig. 2C. The dashed lines indicate the location of the MHB based on information from Fig. 2D. Black arrows indicate the elevated Wnt signaling in the midbrain region adjacent to the MHB. (D) Representative images of *AXIN2* mRNA expression, along with DAPI and OTX2 staining. The bottom images show a zoomed-in view of the colony center. (E) Heatmap plot shows the Pearson correlation coefficient between *AXIN2* and DAPI, and between *AXIN2* and OTX2. As OTX2 protein and *AXIN2* _1_m_0_RNA are found in different cellular compartments, *AXIN2* images were Gaussian filtered with a sigma value of 16.7 pixels, which is approximately one-third of the major axis of the nucleus.

To further investigate these observations, we examined Wnt signaling activity along the colony radius by measuring expression of the direct Wnt target LEF1 alongside the dynamics of AP coordinate acquisition (Fig. 5B–B’). A LEF1 gradient forms by D2 and reshapes over D3–D4. Smaller colonies take longer to stabilize their gradient (Fig. 5B’), initially showing more homogeneous signaling before LEF1 is reduced at the colony center. Comparing across sizes, smaller colonies exhibit higher LEF1 levels and steeper gradients (Fig. S4A). Together, these dynamics likely reflect feedback mechanisms that sharpen the Wnt gradient in smaller colonies. By D4, LEF1 becomes slightly elevated at the colony center, forming a valley-like region of Wnt activity (discussed in the next section).

HOXB1 emerges on D3 with a spatial pattern mirroring LEF1, and higher overall LEF1 corresponds to increased HOXB1 expression (Fig. 5B–B’). This relationship persists on D4, with HOXB1 showing no elevation at the central LEF1 peak—the colony center retains an anterior coordinate. Notably, even the lowest LEF1 level in r50 on D4 exceeds the peak value in other conditions, yet the local dip in LEF1 at the colony center still correlates with reduced HOXB1. This argues against strict thresholding of Wnt activity as the basis for cell fate decisions; instead, cells appear to read differential levels relative to their surroundings. The combination of this feature with the gradient rescaling noted above allows the overall pattern to partially scale with colony size.

### Wnt activity gradient does not increase monotonically along the AP axis

The Wnt activity gradient that rises from anterior to posterior is present in nearly all bilaterally symmetric animals [63], and Wnt activity is generally assumed to vary smoothly along the axis. However, in Fig. 5B’ we observed that LEF1 expression dips on D4 near the MHB in colonies large enough to form both anterior and posterior cell fates. Measuring *AXIN2*, another direct Wnt target, revealed the same trend (Fig. S4B–C, E), indicating that Wnt activity is not monotonic along the axis but is instead elevated in the midbrain relative to both the forebrain anteriorly and the anterior hindbrain posteriorly, producing a local minimum at the MHB itself. This is consistent with earlier observations of local Wnt enrichment in the midbrain in mouse Wnt reporter lines and Xenopus *ω*-catenin staining [35, 36].

To understand how these variations in Wnt signaling map to cell fates, we compared the anterior OTX2-expressing cells with cells also in the central region that lack OTX2, *WNT1*, and the forebrain markers PAX6 and FOXG1, an expression profile suggesting a more posterior fate than the neighboring OTX2-expressing midbrain (Fig. 5D, Fig. 2F, Fig. S1A). The OTX2-expressing area shows higher *AXIN2* levels than the non-OTX2-expressing area (Fig. 5D, Fig. S4F), with a strong correlation between *AXIN2* and OTX2 within the colony center (Fig. 5E). Thus the midbrain territory exhibits higher Wnt signaling than the anterior hindbrain immediately posterior to it. This pattern also holds in the mouse scRNA dataset, where both *Lef1* and *Axin2* show elevated Wnt activity in the midbrain compared to surrounding regions (Fig. 5C), suggesting a relationship between elevated midbrain Wnt activity and MHB formation.

### WNT8A upregulates Wnt activity in the posterior hindbrain and defines the position of the midbrain-hindbrain boundary

The non-monotonic Wnt activity profile suggests that distinct ligands may shape its features in different regions of the axis. Many Wnt ligands are expressed during axial patterning [22–27]. To understand how endogenous Wnt ligands contribute to AP patterning, we first examined the expression dynamics of all Wnt ligands in the mouse scRNA dataset (Fig. 6A’, Fig. S5A) [50]. Several showed AP-dependent expression: *Wnt1*, *Wnt3a*, *Wnt5a*, *Wnt5b*, *Wnt6*, *Wnt7b*, *Wnt8a*, and *Wnt8b*.

**Figure 6:**
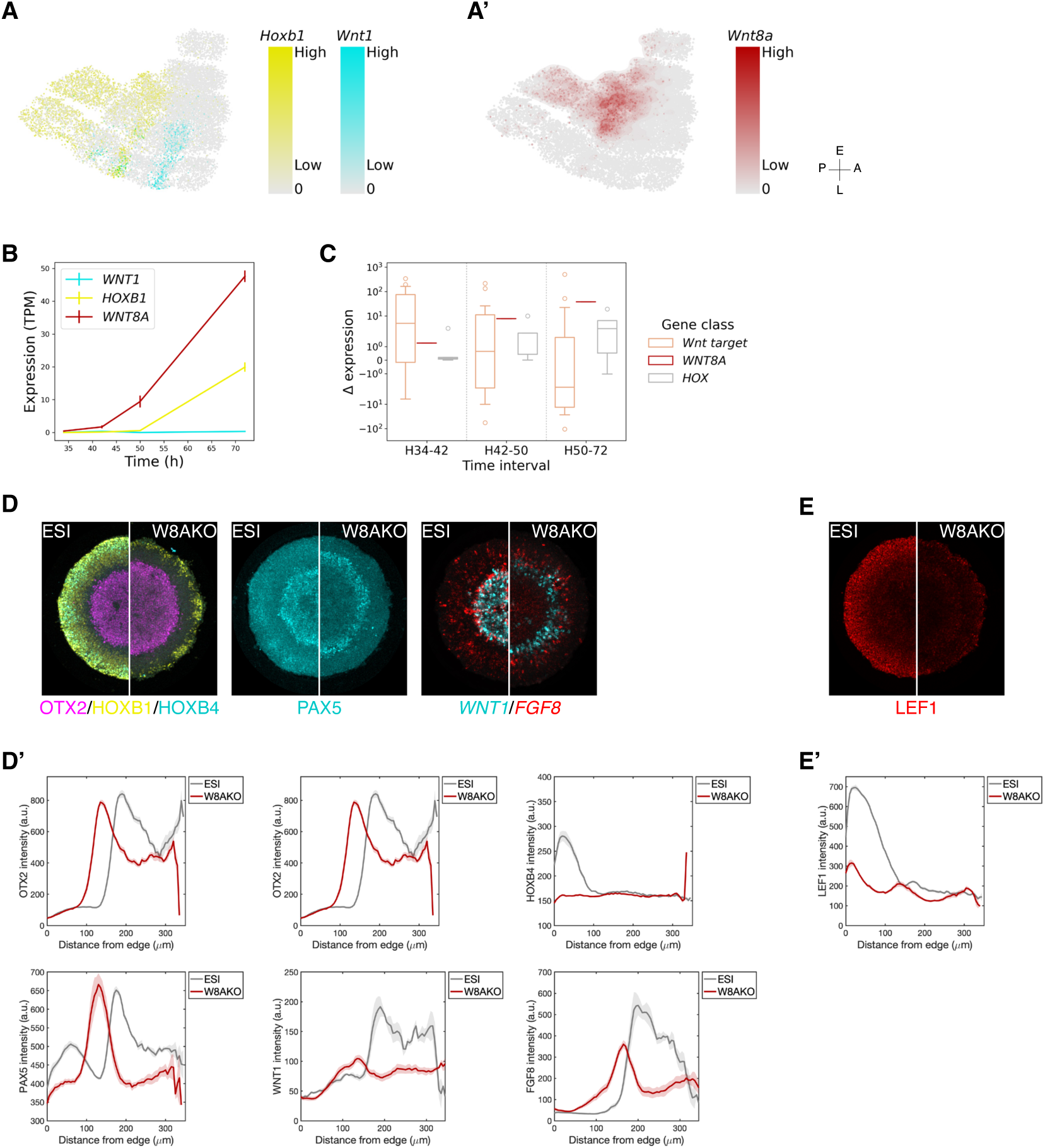
WNT8A is involved in AP coordinate specification. (A, A’) Expression levels of *Hoxb1*, *Wnt1*, and *Wnt8a* overlaid on the UMAP plot shown from Fig. 2A. (B) Quantification of *HOXB1*, *WNT1*, and *WNT8A* from bulk RNA sequencing dataset, averaged across two biological replicates (*N* = 2). (C) Box plot of the expression level change within the specified time intervals. (D, E) Representative immunofluorescence images of indicated markers following the treatments described in Fig. 1C, middle. (D’, E’) Quantification of the expression levels of the indicated markers.

To examine these dynamics in human cells and identify upstream and downstream regulators, we differentiated hPSCs into posterior neural cells in standard culture, adding WNT3A at 34 h as in the micropatterned protocol (Fig. 1C, middle); this treatment is expected to produce cells similar to those near the edge of micropatterned colonies. Samples collected at 34, 42, 50, and 72 h were subjected to bulk RNA sequencing. As anticipated, *HOXB1* was expressed whereas *WNT1* was not, and *WNT8A* was upregulated earlier than *HOXB1* (Fig. 6B), consistent with the mouse expression pattern (Fig. 6A–A’). Following WNT3A addition, direct Wnt target genes including *SP5*, *GBX2*, and *TCF4* were upregulated first, followed by *WNT8A* and then HOX genes (Fig. 6C, Fig. S5B–D). This sequence suggests that *WNT8A* could drive the high posterior Wnt activity and posterior cell fates observed at the colony edge.

To test this directly, we generated WNT8A knockout (KO) cell lines (Fig. S6A–B). During micropatterned differentiation, loss of WNT8A resulted in expansion of the OTX2 domain, reduced HOXB1 and HOXB4 expression, and outward shifts of PAX5, *WNT1*, and *FGF8*, with the colony center filled with PAX6-expressing cells indicating more anterior identities (Fig. 6D–D’, Fig. S6C–D’). Consistent with this anteriorization, peak Wnt activity near the colony edge was strongly reduced (Fig. 6E–E’). Together, these results demonstrate a critical role for WNT8A in specifying hindbrain and spinal cord fates and in positioning the MHB. Even cells at the edge, which respond most strongly to exogenous Wnt, require a self-organized response through endogenous WNT8A to fully posteriorize.

Notably, the Wnt activity profile remained non-monotonic in WNT8A KO colonies, with the local elevation in the midbrain territory preserved. This indicates that the midbrain elevation is independent of WNT8A and likely reflects a different ligand—*WNT1* is a strong candidate given its specific expression in the midbrain both *in vivo* and in our system (Fig. S5A, Fig. 2F).

## 3 Discussion

We developed a system in which human pluripotent stem cells (hPSCs) differentiate to neural precursors and self-organize an anterior-posterior (AP) axis along the radial coordinate of a two-dimensional colony, with cell fates progressing from anterior at the center to posterior at the edge. Neural conversion is induced by dual-SMAD inhibition, after which colonies are treated with WNT3A to confer axial identities. Comparison with published scRNA expression data showed that the system recapitulates both the spatial ordering and the temporal dynamics of gene expression in the developing mammalian nervous system. Using this system, we determined the quantitative relationship between Wnt ligand presentation, the dynamics of signaling activity, and AP patterning. The timing of Wnt addition emerged as a critical parameter: delaying Wnt allowed central cells to commit to anterior fates while edge cells were posteriorized, with later addition producing larger anterior regions. The self-organizing nature of the system distinguishes it from recent models that impose an AP axis using microfluidic Wnt gradients [39, 40], and the resulting patterns of endogenous Wnt activity prove more complex than a simple monotonic gradient, as discussed below.

Multiple models have been proposed to explain how the pluripotent cells of the epiblast both convert to neural fate and acquire their position along the AP axis. The classic model holds that cells are first specified to an anterior neural identity, after which cells outside the forebrain are posteriorized according to their position along the axis [12, 17, 64]. Emerging evidence challenges this view, suggesting that hindbrain and spinal cord are specified as posterior before acquiring neural identity [14–16]. Our data support the latter view: AP position markers such as OTX2 and HOXB1 are expressed concurrently with pluripotency-associated genes such as OCT4 and ECAD, and well before the emergence of neural fate markers such as SOX1 or PAX6 (Fig. 2B, E–E’, Fig. S1).

The role of Wnt signaling in establishing the AP axis is remarkably conserved across the animal kingdom, involving posterior expression of Wnt ligands and anterior expression of Wnt inhibitors, which together generate a Wnt activity gradient [19, 63]. Measurements in our system, however, reveal that the Wnt profile does not simply increase along the axis. Instead, Wnt activity is elevated in the midbrain relative to adjacent regions, creating a local dip in the overall gradient that aligns with the anterior hindbrain. By examining scRNA expression data from the mouse embryo, we found that Wnt target expression follows a similar pattern along the AP axis (Fig. 5B–E), suggesting that this feature is not peculiar to our *in vitro* system. Many hypotheses have been proposed for how the midbrain-hindbrain boundary (MHB) is established and maintained [65]. In our system, the local elevation of Wnt activity in the midbrain emerges after the OTX2/HOXB1 boundary is already established, indicating that this feature is unlikely to be required for initial MHB specification. It may instead contribute to maintaining the boundary, or reflect feedback between the developing Wnt profile and the cell fates it has helped specify. Testing these possibilities will require spatial manipulation of Wnt signaling to alter its profile along the axis; for example, elevating Wnt activity in the anterior hindbrain to match midbrain levels would test whether the MHB can still be maintained without a local reversal of the gradient. Among the 19 Wnt genes in the mouse genome, seven (*Wnt2b*, *Wnt3*, *Wnt3a*, *Wnt5a*, *Wnt5b*, *Wnt8a*, and *Wnt11*) are expressed in the primitive streak. Four Wnt genes show strong AP-related phenotypes upon knockout (KO): *Wnt1* KO has midand hindbrain deficiencies, *Wnt3* KO results in failure of AP axis formation prior to gastrulation, and *Wnt3a* and *Wnt5a* KOs have truncated AP axes [1, 66]. *Wnt8a* KO mice are viable and show no obvious phenotype on their own, with *Wnt8a* required for normal anterior trunk development only in the absence of *Wnt3a* [66, 67]. In standard hPSC culture, overexpression of *WNT8A* produces a weaker posteriorizing effect than *WNT3A* [68], indicating that *WNT8A* alone is not sufficient to drive strong posteriorization. Our results show that it is nonetheless necessary: *WNT8A* KO in our system caused loss of hindbrain and spinal cord fates despite the presence of exogenous *WNT3A*, indicating that endogenous *WNT8A* cannot be fully replaced by exogenous *WNT3A* even when the latter is supplied at posteriorizing levels. The contrast with the mild mouse *Wnt8a* KO phenotype could reflect a species difference in Wnt ligand requirements, but could equally arise from compensation by Wnt ligands produced by neighboring tissues *in vivo* — sources that are absent from the isolated micropatterned colony. Our system could be used to dissect the roles of individual Wnt ligands at distinct AP coordinates in human neural development; *WNT1*, *WNT5A*, and *WNT10B* are all interesting candidates for future investigation (Fig. S5C).

In this study, we primarily focused on generating patterns spanning anterior (OTX2-expressing) to posterior (HOX gene-expressing) fates. Under WNT3A treatment, the system does not form the most anterior forebrain (FOXG1expressing) or the most posterior spinal cord fates, but the range it does access is tunable: adjusting the timing, level, or duration of WNT3A treatment shifts the system along the AP axis. Replacing WNT3A with the PORCN inhibitor IWP2 allows the colony to reach the most anterior telencephalic fates (Fig. S1C–C’), further broadening the system’s applicability. The failure to generate the most posterior spinal cord fates likely reflects the absence of two requirements: the generation and maintenance of neuromesodermal progenitors (NMPs), which give rise to the more caudal spinal cord, and the longer culture times needed for full posteriorization [14, 15, 69–71]. Adding FGF signaling might in principle sustain NMPs at the colony edge and allow further posteriorization before neural differentiation; however, our recent work on micropatterns found that combined Wnt and FGF treatment instead drove differentiation toward endoderm or paraxial mesoderm depending on the state of the Nodal pathway [72]. Optimizing the concentrations and timing of Wnt and FGF, potentially combined with inhibitors of mesendodermal fates, may identify conditions that stably maintain NMPs at the colony edge while allowing the central cells to undergo neural differentiation.

Micropatterned colonies self-organize into patterns of cell fates in a variety of contexts, including germ layer patterning, medial-lateral patterning of the ectoderm, and the AP patterns described here (Fig. 1) [41, 42, 46, 73]. In all of these systems, exogenous signals are most strongly received at the colony edge, which then acts as a signaling center, initiating cascades of endogenous signals that generate the pattern [48]. This organization may mimic tissue boundaries in the embryo, such as the boundary between the amnion and epiblast during germ layer patterning. A limitation, however, is that the colony has only a single signaling center, which prevents the system from modeling the entire AP axis at once — full coverage would require both Wnt activators and inhibitors acting from distinct sources. Juxtaposing cell populations expressing different signaling molecules, or using optogenetic tools to control signaling in space, are promising approaches for systems with multiple signaling sources [74, 75]. For example, neural-differentiating cells could be placed between two populations, one expressing Wnt activators and the other Wnt inhibitors. Alternatively, the AP patterning system developed here could be combined with optogenetic induction of Wnt inhibitors at the colony center, allowing a more complete recreation of the AP axis.

Although our system captures only part of the AP axis at a time, the tunability of Wnt parameters and the ability to combine it with the strategies above provide a foundation for more complete models of AP patterning. The system has allowed us to dissect how Wnt signaling shapes pattern generation and to uncover features such as the non-monotonic Wnt profile and the requirement for endogenous WNT8A. Future work using this system and its variants can explore how Wnt interacts with other pathways such as FGF and retinoic acid to fully pattern the axis.

## 4 Materials and Methods

### Cell lines

Experiments were performed using ESI017 hESCs (obtained from ESI BIO, RRID: CVCL_B854) unless otherwise noted. For live imaging of WNT signaling dynamics, the H9 TCF/LEF:H2B-GFP reporter cell line was used [62].

### Generation of knockout cell lines

To generate WNT8A knockout (KO) hESC lines, we used CRISPR-Cas9 genome editing with ribonucleoprotein (RNP) complexes delivered via electroporation. Guide RNAs (gRNAs) were designed using the Synthego CRISPR Design Tool (spCas9, H. sapiens) and two non-overlapping gRNAs were selected that target early exons. Synthetic sgRNAs (Synthego) were complexed with recombinant Cas9 protein (IDT, Alt-R S.p. Cas9) at a 1:2 ratio (1.2 µM Cas9: 2.4 µM sgRNA) at 37*^→^*C for 15 min.

hESCs were dissociated with Accutase, washed in PBS--, and resuspended in P3 Nucleofector^TM^ Solution (Lonza). RNPs and 220,000 cells were electroporated in 20 µL total volume using the 4D-Nucleofector^TM^ X Unit. Postelectroporation, cells were incubated for 10 min with 80 µL of pre-warmed medium and then plated in 12-well plates. Medium was refreshed daily.

Genomic DNA from bulk-edited cells was isolated at 72 h (QIAwave DNA Kit, Qiagen), and target loci were PCR-amplified and analyzed via Sanger sequencing and TIDE deconvolution. Populations with >70% indels were clonally isolated by flow cytometry and validated by TOPO TA cloning (Invitrogen) and Sanger sequencing. Biallelic KO clones were expanded for downstream experiments.

### Routine cell culture

All cells were grown in the chemically defined medium mTeSR1 (STEMCELL Technologies) and kept at 37*^→^*C, 5% CO_2_. Cell culture dishes were coated with Matrigel (Corning) or Geltrex (Thermo Fisher Scientific), each 1:500 diluted in PBS-and incubated at 4*^→^*C overnight. Cells were regularly checked for pluripotency and mycoplasma contamination throughout the study. Dispase (STEMCELL Technologies; 1:5 dilution in DMEM/F12 (VWR)) was used for gentle dissociation and routine passage. Accutase (Corning) was used to prepare single-cell suspensions. ROCKinhibitor Y-27632 (STEMCELL Technologies; 10 µM, diluted 1:1000) was used to reduce dissociation-induced apoptosis during single-cell suspension.

### Micropatterning

Micropatterning experiments were performed on either micropatterned chips or 96-well micropatterned plates from CYTOO.

For chips, hESCs were seeded onto micropatterned surfaces coated with 5 µg/mL LN-521 (Biolamina; diluted in PBS++). The modified mTeSR1 protocol from [76] was used with the following modifications: 1) the chip was incubated for 3 h at 37*^→^*C during coating; 2) the chip was washed eight times with PBS++ (cytiva); 3) 1 *→* 10^6^ cells were deposited onto the chip using a circular motion; 4) seeded cells were incubated for 1.5 h at 37*^→^*C, followed by two partial washes and one full wash (4 mL pre-warmed PBS-in each wash).

For 96-well plates, hESCs were seeded onto micropatterned surfaces coated with 50 µL of 5 µg/mL LN-521 and incubated for 2.5 h at 37*^→^*C or overnight at 4*^→^*C. The LN-521 was removed by three partial washes and one full wash (100 µL PBS++ in each wash, except the first wash which was 150 µL). The plate was either used immediately or stored for up to 2 weeks at 4*^→^*C with PBS++ covering the surfaces. For seeding, cells were washed twice with PBS-and removed from the surface using Accutase (Corning) for 5–7 min at 37*^→^*C. Subsequently, cells were harvested by centrifugation (1000 rpm or 300 g for 4 min) and re-suspended in mTeSR1 containing ROCKi. 120,000 cells were then seeded into each LN521-coated well and incubated for 30 min at 37*^→^*C. To remove non-specifically bound cells in the uncoated region, cells were washed once with PBS--, then switched to mTeSR1 and incubated for 1 h at 37*^→^*C (no ROCKi contained). Treatment cocktails described in the text were then added.

### Differentiation

Neural induction was performed in N2B27 medium supplemented with SB431542 (10 µM; Fisher Scientific) and LDN193189 (100 nM; Fisher Scientific). N2B27 was prepared by filtering a mixture of 24 mL DMEM/F12 (Corning), 24 mL Neurobasal media (Life Technologies), 0.5 mL N2 supplement (Fisher Scientific), 1 mL B27 supplement without vitamin A (Fisher Scientific), 0.5 mL Glutamax (Gibco) and 50 µL *ω*-mercaptoethanol (Gibco). In all experiments, the medium was replenished daily. However, when Wnt was added at D1.5, the medium was first replenished at D1.5 and then again at D3.

### Immunostaining

Cells were fixed in 4% paraformaldehyde (Electron Microscopy Science) for 10 min at room temperature, followed by one PBS-wash. Fixed samples were then permeabilized for 30 min at room temperature with blocking buffer (PBS-with 0.1% Triton X-100 (Sigma-Aldrich) and 3% donkey serum (Sigma-Aldrich)). Primary antibodies were diluted in blocking buffer as listed in Tbl. 1 and applied to samples overnight at 4*^→^*C (this is recommended for better staining rather than 2 h at room temperature). Samples were then sequentially washed three times with PBST (PBS with 0.1% Tween 20 (Sigma-Aldrich), 10 min each time), incubated for 30 min at room temperature with secondary antibodies (Tbl. 1) supplemented with DAPI (1:1000), and washed three times with PBST in the same manner as before. For chips, coverslips were then mounted in Fluoromount-G (Southern Biotech) and allowed to dry in the dark for several hours. For the 96-well plate, samples were kept in PBS-and were ready for imaging.

### Single-molecule fluorescence *in situ* hybridization

Cells were fixed in 4% paraformaldehyde for 10 min at room temperature. smFISH was performed following the manufacturer’s instructions (ACD Bio, RNAscope Multiplex Fluorescent v2 Assay). Probes are listed in Tbl. 1.

### Imaging

#### Fixed cell imaging

All images were acquired at 10! (NA 0.40, zoom 1.7!) and 30! (NA 1.05, zoom 2!) using an Olympus FV1200 laser scanning confocal microscope (LSM) with FV10-ASW 4.2 software.

#### Live cell imaging

TCF/LEF1:H2B-GFP H9 cells were seeded in multiple wells of a 96-well plate and differentiated as described above. Five colonies (700 µm) per well were imaged using a 20! (NA 0.75) objective on Olympus/Andor spinning disk confocal microscope. Every 1 h, multiple z-planes were acquired per position and three positions were set to cover one colony. Imaging was halted for a short period to replenish media at the exact replenishing time in the fixed cell experiment. Cells were maintained in a chamber at 37*^→^*C and 5% CO_2_ throughout the duration of imaging.

### Image analysis

All experiments were repeated at least twice with consistent results. ‘n’ in the figure captions denotes the number of micropatterned colonies imaged per condition within the same experiment. Unless otherwise stated, *n* = 5. Colonies were selected solely based on images from the brightfield or DAPI channel without considering other channels to avoid observer bias.

Images were batch-stitched using FIJI with a customized Python script ([77]). Other analyses were performed using customized MATLAB code. Both are available at https://github.com/warmflashlab/Du2025Novel. Micropatterned colonies were automatically identified, after which the edge or center of each colony was defined, and a specified marker profile was quantified as a function of intensity versus distance to the edge/center. Distance was binned in 5 µm increments, estimated to be the radius of a cell. For each condition, the mean intensity at each distance was averaged across all the colonies, and error bars represent the standard error of the mean intensity.

### Bulk RNA sequencing

#### Sample collection and quality control

hESCs were seeded at the density of 30,000 per cm^2^ in a 24-well plate with mTeSR1 and ROCKi 24 h before treatment. hESCs were under neural induction (described in Sec. 4) and treated with WNT3A (300 ng/mL) at 34 h. Samples were collected at 34, 42, 50, 72 h. Total RNA was extracted using RNAqueous-Micro Total RNA Isolation Kit (Invitrogen) and stored at *↑*80*^→^*C. RNA integrity was checked by Nanodrop, agarose gel electrophoresis, qPCR, and the Agilent 2100 system. Sequencing was performed by Novogene Co. using the Illumina paired-end 150 platform (NovaSeq X Plus). Another biological repeat was performed using the same protocol.

#### Analysis

To quantify the abundance of transcripts for each sample, Salmon (v1.9.0) was used with the selective alignment method. Reads were aligned to the human GRCh38 transcriptome using the precomputed salmon_sa_index:default. We quantified transcripts with Salmon using the `-l A` flag to infer the library type (paired-end) automatically and the `--validateMappings` flag to use the selective alignment method. The resulting datasets were then processed using the tximeta package in R, and the counts and abundance matrices were exported for downstream analyses. Both the processed data and code are available at https://github.com/warmflashlab/Du2025Novel. The complete datasets are deposited in Gene Expression Omnibus (GEO) repository with accession number GSE337980.

### Single-cell RNA sequencing analysis

The mouse single-cell RNA-sequencing dataset was acquired from [50] and was normalized using Scanpy and scVItools ([51, 78]). A UMAP plot of all cell fates was generated to compare with the original plot and verify the reproducibility of our normalization. Only cell fates relevant to our study were selected: rostral neuroectoderm, caudal neuroectoderm, forebrain/midbrain/hindbrain, spinal cord, neuro-mesodermal progenitor (NMP). These cell types and stages were plotted again in UMAP to assist in the interpretation of the spatial and temporal expression of the genes of interest. The normalized data and code are available at https://github.com/warmflashlab/Du2025Novel.

## Acknowledgments

We would like to thank Zachary Kingston for programming advice and Uny Tian for assistance with FISH staining. This work was funded by National Institutes of Health awards R01GM126122 and R35GM149328.

