## Supplementary material for "A novel self-organizing human pluripotent stem cell system reveals the role of Wnt signaling parameters in anterior-posterior patterning of the nervous system": Figures S1-S5, Table S1

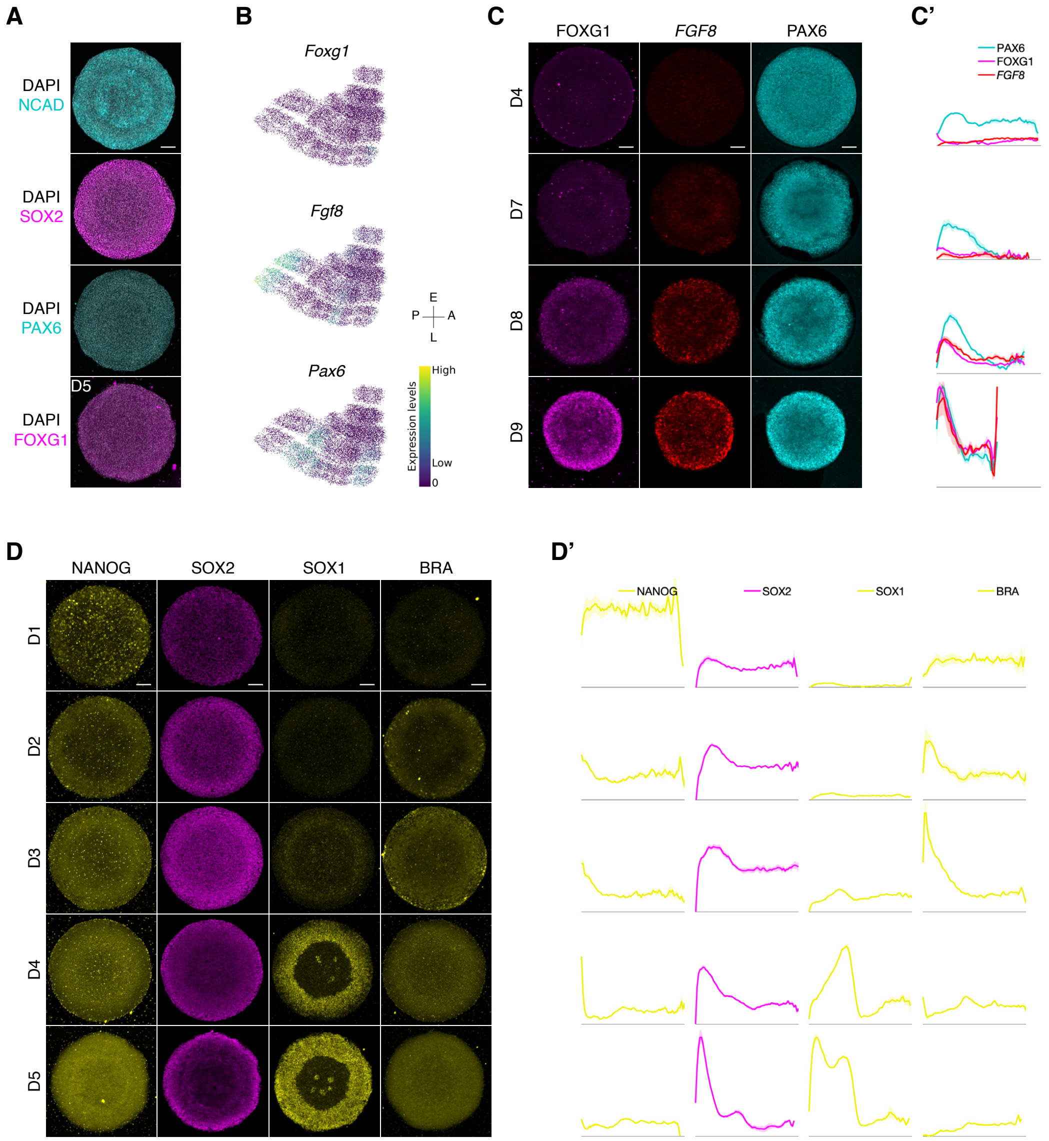


**Figure S1: Dynamics of anterior and posterior marker expression.** (A) Representative immunofluorescence images of the indicated markers. WNT3A was added at D1.5 during the 5-day neural induction. All markers were observed at D4 except for FOXG1. (B) Expression levels of the indicated genes overlaid on the UMAP plot shown in Fig. 2A. (C, D) Representative immunofluorescence images of the indicated markers from D4–D9 and D1–D5 respectively. *FGF8* is mRNA expression. (C', D') Quantification of the expression levels of the indicated markers as a function of distance from the colony edge on each indicated day.

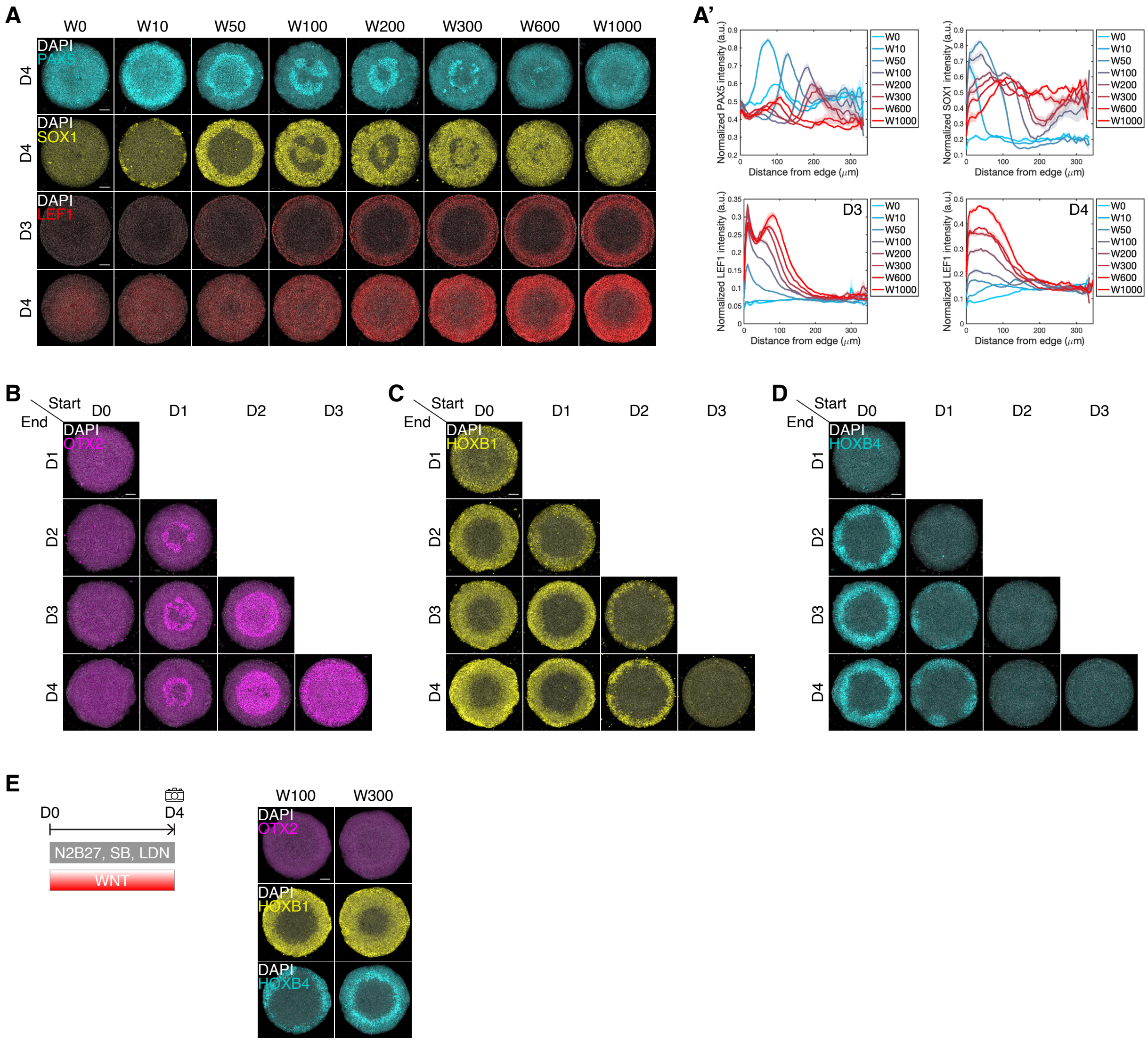


**Figure S2: AP patterns under Wnt treatments with varying concentrations, start and end times.** (A) Representative immunofluorescence images of the indicated markers. Different concentrations of WNT3A $(\mathrm{ng/mL})$, as specified in the labels, were added on D1.5. (A') Quantification of the expression levels of LEF1, PAX5, and SOX1 following the treatments described in A, normalized to DAPI intensity. (B, C, D) Representative immunofluorescence images of OTX2, HOXB1 and HOXB4, arranged in a checkerboard layout, with the WNT3A addition and removal times indicated along the horizontal and vertical axes. (E) Representative immunofluorescence images of the indicated markers. The concentrations of WNT3A $(\mathrm{ng/mL})$, as specified in the labels, were added on D0.

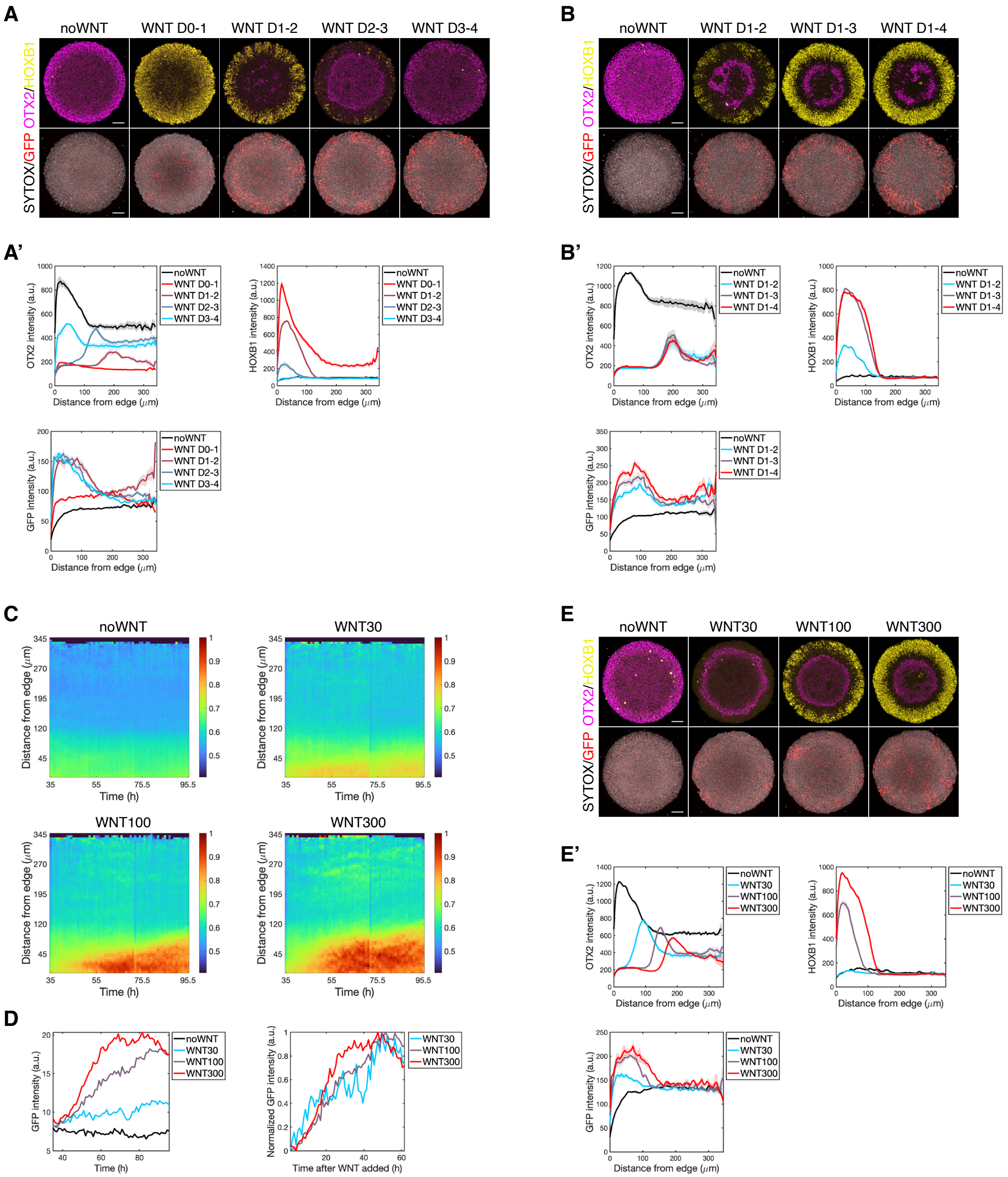


**Figure S3: Wnt signaling dynamics under different WNT3A concentrations.** (A, B, E) Representative immunofluorescence images of the indicated markers, following 4-day neural induction and WNT3A treatment as described in the labels. (A', B', E') Quantification of the expression levels of OTX2, HOXB1 and GFP. (C) Heatmap plots of GFP levels, with the x-axis representing experimental time and the y-axis representing the distance from the colony edge, following 4-day neural induction in which WNT3A was added at D1.5 with the concentration (ng/mL) specified in the labels. (D) Quantification of average GFP intensity in the ring-shaped domain (0–100 μm from the edge) as a function of time. Normalized GFP intensity is calculated by scaling the GFP levels to their starting (base) and peak (maximum) values.

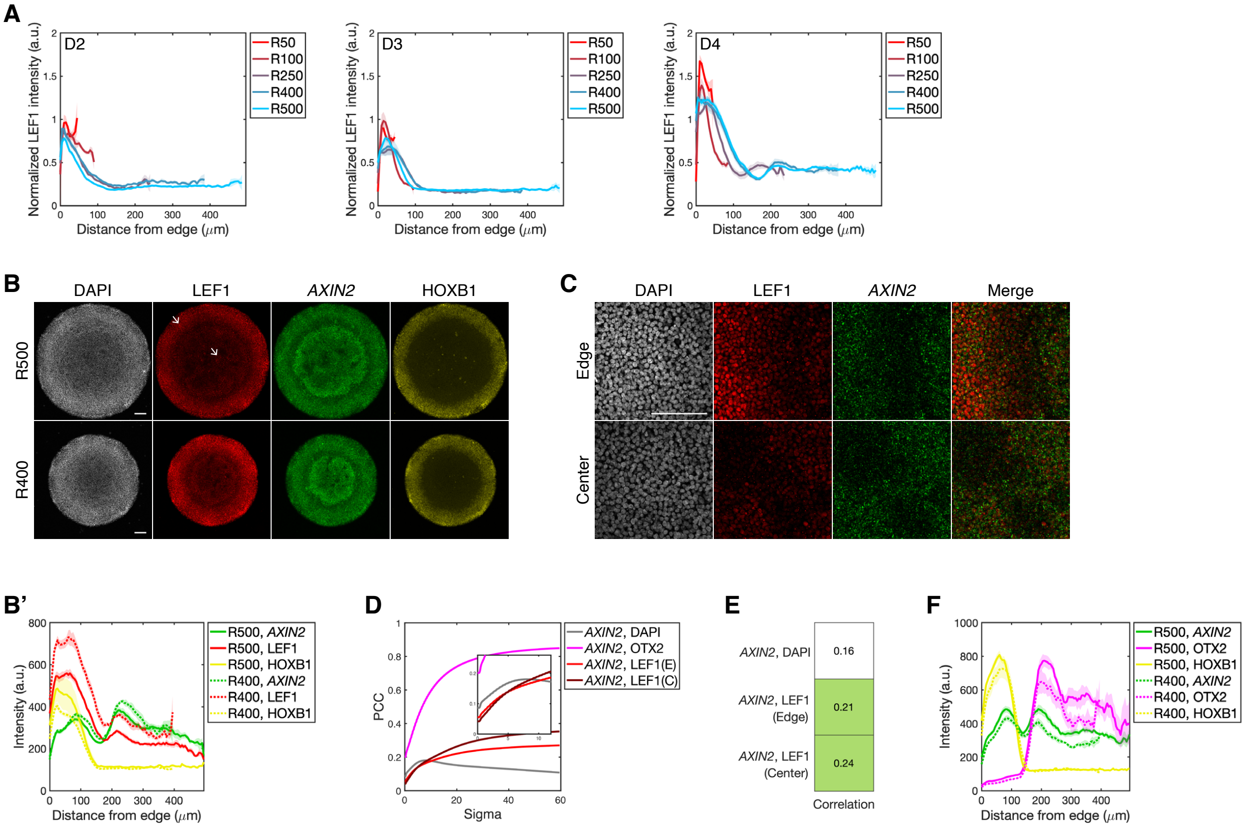


**Figure S4: LEF1 and *AXIN2* show the same expression trend along the AP axis.** (A) Quantification of LEF1 expression levels, normalized to DAPI intensity. For the indicated colony radii, $n = 5$ for R500, R400 and R250; $n = 8$ for R100; and $n = 9$ for R50. (B) Representative images of *AXIN2* mRNA expression, along with immunofluorescence for DAPI, LEF1 and HOXB1. White arrows indicate the edge and center of the colony. (B') Quantification of the expression levels of *AXIN2*, LEF1 and HOXB1. (C) Representative images of *AXIN2* mRNA expression, along with immunofluorescence for LEF1 at the edge and center of the colony. The colony diameter is 1000 $\mu$m. (D) *AXIN2* images were Gaussian filtered with sigma values ranging from 0–60. PCC refers to the Pearson correlation coefficient. The inset plot shows the sigma range from 0–12 in greater detail. (E) Heatmap plot shows the Pearson correlation coefficient between the indicated pairs. *AXIN2* images were Gaussian filtered with a sigma value of 16.7. (F) Quantification of the expression levels of *AXIN2*, OTX2 and HOXB1.

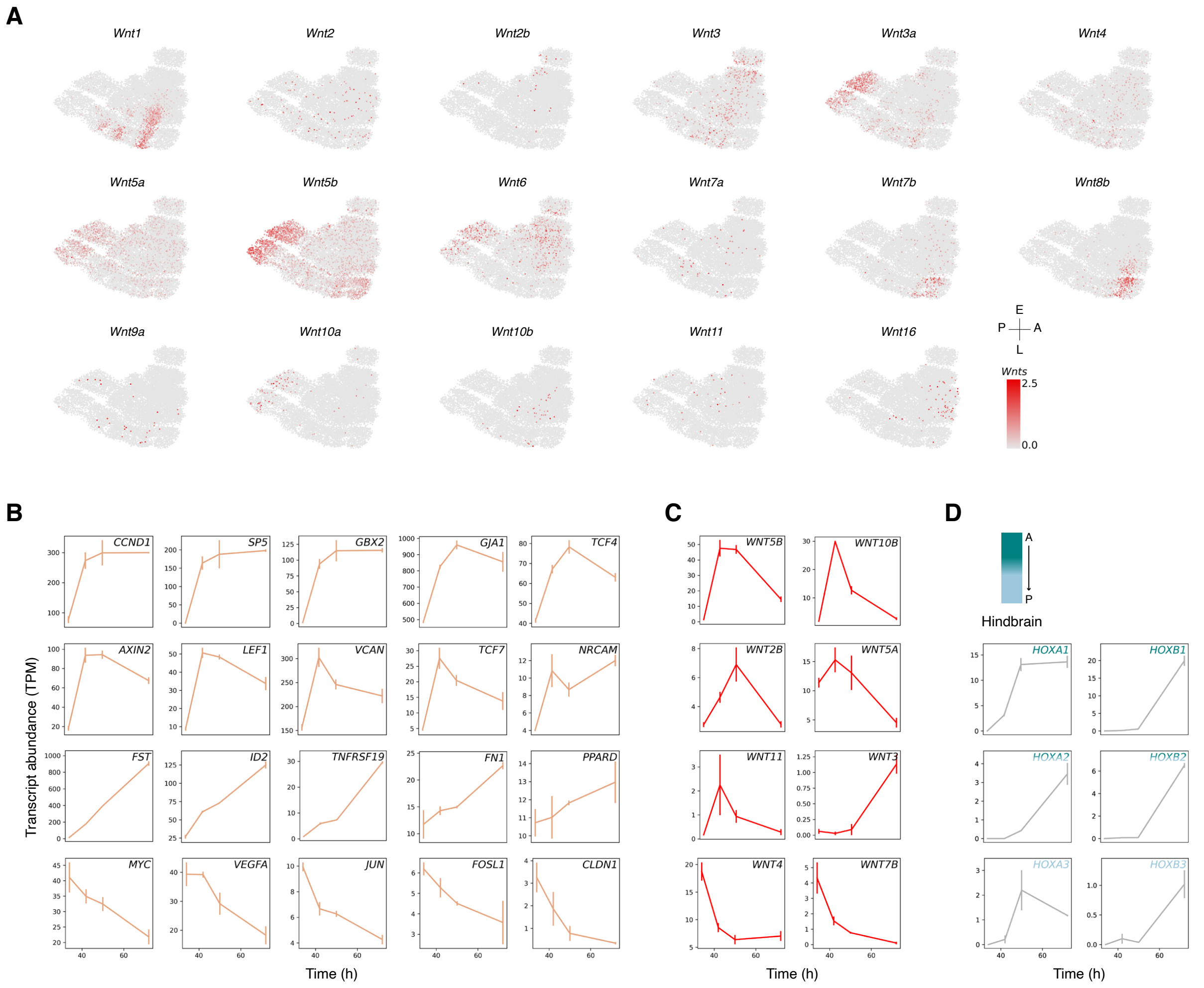


**Figure S5: Wnt ligands and related genes expression pattern in mouse and human.** (A) Expression levels of all Wnt ligands other than *Wnt8a* overlaid on the UMAP plot shown from Fig. 2A. Dynamics of expressed (B) Wnt direct target genes, (C) Wnt ligands, and (D) HOX genes, with expression levels averaged across two biological replicates ($N = 2$). Direct Wnt targets are defined as genes with validated Tcf/Lef binding sites for which those sites have been shown to be required for Wnt-dependent expression (`https://wnt.stanford.edu/target_genes`). In D, anterior hindbrain *HOX* genes are shown in teal and posterior hindbrain *HOX* genes in light blue.

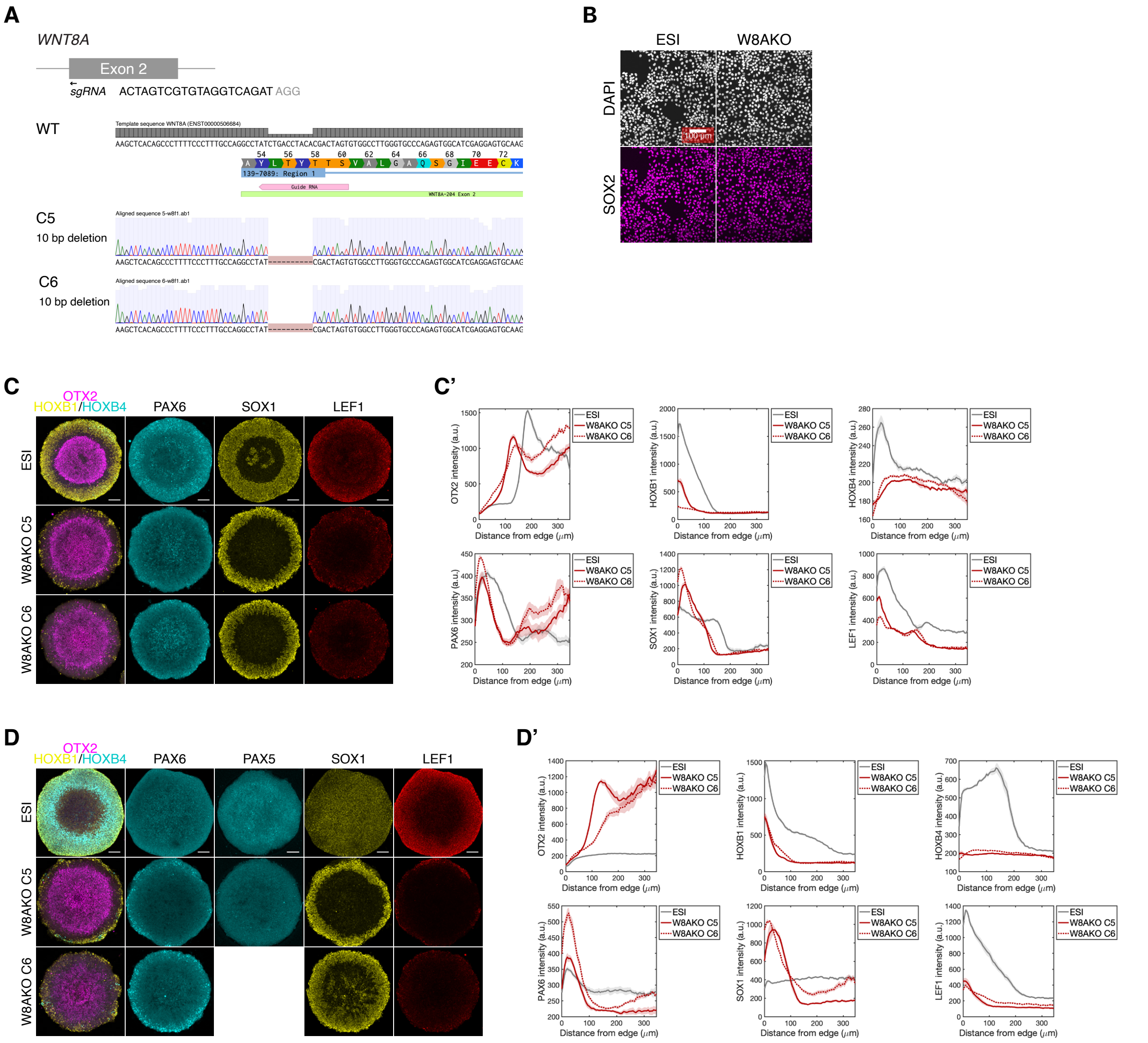

**Figure S6: Creation and validation of WNT8A knockout cells.** (A) sgRNA is used to make a double-stranded break on exon 2 of *WNT8A*. Genomic sequence of *WNT8A* locus in corresponding knockout cells. (B) Representative immunofluorescence images of DAPI and pluripotency marker SOX2. (C, D) Representative immunofluorescence images of indicated markers following the treatments described in (C) Fig. 1C, middle and (D) Fig. 1B. (C', D') Quantification of the expression levels of the indicated markers. The colony diameter in C–D is 700 µm. Scale bar, 100 µm.

| Antibody | Vendor | Catalog number | Dilution | Species |
|---|---|---|---|---|
| OTX2 | Abcam | ab322111 | 1:500 | Rabbit |
| HOXB1 | R&D Systems | AF6318 | 1:200 | Sheep |
| HOXB4 | DSHB | I12 | 1:100 | Rat |
| OCT4 | BD Biosciences | 611203 | 1:400 | Mouse |
| ECAD | Cell Signaling Technology | 3195S | 1:200 | Rabbit |
| NCAD | Sigma-Aldrich | C2542 | 1:300 | Mouse |
| CDX2 | Cell Signaling Technology | 12306s | 1:100 | Rabbit |
| PAX5 | DSHB | PCRP-PAX5-1B1 | 1:20 | Mouse |
| CTNNB1 | R&D Systems | AF1329 | 1:200 | Goat |
| LEF1 | Cell Signaling Technology | 2230s | 1:200 | Rabbit |
| SOX2 | Cell Signaling Technology | 23064s | 1:200 | Rabbit |
| FOXG1 | Abcam | ab18259 | 1:500 | Rabbit |
| PAX6 | DSHB | PAX6 | 1:50 | Mouse |
| NANOG | R&D Systems | AF1997 | 1:200 | Goat |
| SOX1 | R&D Systems | AF3369 | 1:100 | Goat |
| Alexa 405 anti-Rabbit | Abcam | ab175651 | 1:400 | Donkey |
| Alexa Fluor 488 anti-Rat | Life Technologies | A21208 | 1:400 | Donkey |
| Alexa Fluor 488 anti-Mouse | Life Technologies | A21202 | 1:400 | Donkey |
| Alexa Fluor 555 anti-Rabbit | Life Technologies | A31572 | 1:400 | Donkey |
| Alexa Fluor 647 anti-Goat | Life Technologies | A11055 | 1:400 | Donkey |
| Alexa Fluor 647 anti-Sheep | Life Technologies | A21448 | 1:400 | Donkey |
| **Probe** | **Vendor** | **Catalog number** | **Dilution** | **Channel** |
| *WNT1* | ACD | 429421 | NA | C1 |
| *FGF8* | ACD | 415791-C2 | 1:50 | C2 |
| *AXIN2* | ACD | 400241-C3 | 1:50 | C3 |

**Table 1: Antibody and probe information.**
